# Tracking animal routes in 3D space through reconstructed habitats from dynamic videos

**DOI:** 10.64898/2026.07.22.740183

**Authors:** Martha M. M. Daniel, Matteo Santon, Ajay Narendra, Martin J. How

## Abstract

Contextualizing the movements of animals into their three-dimensional (3D) habitat contexts is still a major challenge for fields relying on animal tracking methods. This promises to change as Structure-from-Motion photogrammetry tools and techniques revolutionize image processing into the truly 3D spatial realm, by enabling reconstructions of habitat models from overlapping photographs. Combined with tracking data, these techniques would help elucidate drivers behind animal movements that have been previously masked by two-dimensional approaches. Unfortunately, tracking methods are often still impractical for use with understudied or non-model animals, especially those living underwater. In this paper, we describe a method for tracking the translational movements of animals into a photogrammetric habitat model. Our approach spans three general parts: (1) filming the navigation paths of wild animals by following individuals with small cameras (GoPros) on extendable sticks whilst SCUBA diving, (2) reconstructing a 3D spatial habitat model from separate footage, and (3) manually plotting the 3D trajectories of animals into the habitat model. We used one popular commercial software for photogrammetric reconstruction, trajectory plotting, and measurement of trajectories, after which the plots can be exported in a variety of formats for further analyses. Straightforward and flexible methodologies such as this stand to encourage more fieldwork concerning animals that live in structurally complex habitats, or animals that are underrepresented in movement or navigation research. We expect that this approach can be adapted to study many aquatic or terrestrial animals in different habitats, and at various scales.

## Introduction

The movement ecology framework proposed by Nathan et al. (2008), and drawing inspiration from Aristotle’s *De Motu Animalium*, is an effort to integrate different scientific disciplines that share an interest in organismal movements. This paradigm addresses four common challenges for the analysis of organismal movement data, namely the need to define the movement steps, phases, and phenomena that make up an organisms’ path across spatiotemporal scales, and to situate these in environmental context. The movement ecology framework centers on four components, referred to as an organism’s internal state, motion capacity, and navigation capacity, and the external information available to the organism. External information is so intimately linked to navigation capacity that these two components are necessarily defined in terms of each other, yet there seems to remain an unfortunate disconnect between the fields of movement ecology (e.g. Nathan et al. 2008) and sensory ecology (e.g. Dangles et al. 2009; Willemart 2023). Perhaps this is because the historic perspectives of these fields are rooted at opposite extremes of scale (i.e. bird migration behavior: as mentioned by Börger 2016, and sensory physiology: Land 1980 and Dangles et al. 2009). Demšar et al. (2015) conclude that movement research faces five remaining challenges, the first of which concerns navigation and implies that behavioral experiments, sensory perception mechanisms, and neurobiological research are deficient or missing in animal navigation research. Fortunately, these aspects have been linked in sensory ecology definitions at least since 1992 (e.g. Dangles et al. 2009; Willemart 2023), preceded by more than a century of sensory orientation and navigation research (e.g. Turner 1907; Macgregor and Thorpe 1948; Ritz et al. 2000) that is ongoing (e.g. Ritz et al. 2000; Lohmann et al. 2012; Patel and Cronin 2020; Dreyer et al. 2025). Most recently, a collection of universal navigation principles have been proposed (Wirth et al. 2026), including navigational phases and state transitions based on five modules that encompass the same drivers described by the components of movement ecology (Nathan et al. 2008). Interdisciplinary efforts, similar to MOVE (Demšar et al. 2015), would surely benefit both sides in future.

Regardless of discipline, the current methods concerned with movements in space and time tend to fall under the broad classification of “tracking.” Tracking can refer to telemetry methods, which range from locating animals via acoustic transmitters (e.g. Donaldson et al. 2014; Matley et al. 2023), to following migrations via satellite radio telemetry (e.g. Wikelski et al. 2007), to monitoring physiological variables via biotelemetry (e.g. Cook et al. 2004), and to biologging accelerations (e.g. Penna et al. 2015) or environmental measurements along animal routes (e.g. Rutz and Hays 2009). Tracking may also refer to tracing local translational movements in fixed frame videos (e.g. Hedrick 2008; Panadeiro et al. 2021; Chiara and Kim 2023), to estimating fine-scale poses or gaits by tracking body parts in videos (e.g. Mathis et al. 2018; Sin-Hang Yeung et al. 2026), to classifying behaviors (e.g. Troscianko et al. 2026), or to following eye movements (e.g. Land et al. 1990; Billington et al. 2020). Our paper comes from the video-centric side of tracking.

The tracking of animals has allowed researchers to conceptualize drivers behind foraging, migration, and behavioral compensation (e.g. Beltran et al. 2024), as well as characteristics of life history, reproduction, survival, physiology, and spatial use (e.g. Matley et al. 2023), and it is helping researchers understand feedback loops that either hold ecosystems together or induce change (e.g. Russo et al. 2023). By tracking animals, researchers have also explained many aspects of navigation (e.g. Wirth et al. 2026) and teased apart the saliences of senses for certain goals and conditions (e.g. Macgregor and Thorpe 1948; Wolf and Wehner 2000). Of course, there is much left to be discovered about movement in the wild, and interdisciplinary exchanges or collaborations could extend perspectives and tools to address the long-standing limitations within respective disciplines (e.g. Dangles et al. 2009; Munoz and Blumstein 2012; Demšar et al. 2015; Dominoni et al. 2020; Elmer et al. 2021; Hrncir et al. 2023; Williams et al. 2023).

Tracking methods all have limitations in terms of resolution, duration, cost, and/or practicality, (e.g. Donaldson et al. 2014; Kays et al. 2015; Panadeiro et al. 2021; Troscianko et al. 2026), but a common limitation is that most tracking research has, until recently, accounted for movement in just two spatial dimensions (e.g. Tracey et al. 2014; Kays et al. 2015; Russo et al 2023; Lennox et al. 2025). The same can be said for modelling the external information available to animals in their habitats (e.g. Murray and Zeil 2017; Lepczyk et al. 2021; Lennox et al. 2025). It follows that, fully contextualizing the movements of animals into environmental contexts is still a major challenge for movement research (e.g. Matley et al. 2023; Lennox et al. 2025), as is the continued oversight or exclusion of certain types of animals and habitats from study (e.g. Beltran et al. 2024; Elmar et al. 2021; Tekam et al. 2026). Perhaps by prioritizing this challenge of spatial and environmental contextualization (Hrncir et al. 2023; Williams et al. 2023), and by diversifying the habitat types/scales targeted, we would be better equipped to address the other challenges identified for movement research.

Such a priority would benefit from the concepts and tools offered by landscape and seascape ecology. This field concerns the study of relationships between organisms and the physical geometry or spatial patterns of their surroundings (e.g. Boström et al. 2011; Lepczyk et al. 2021; Russo et al. 2023; Remmers et al. 2024). Its research questions usually operate at broader spatial scales, so data collection now relies on various remote sensing methods, including both airborne lidar and shipborne sonar sensors (e.g. Costa et al. 2009; Brock et al. 2009). Fusing such data sources with multispectral or hyperspectral imaging has opened many possibilities for ecologically relevant analyses (e.g. Lepczyk et al. 2021).

While the concepts in landscape and seascape ecology developed mostly from 2D analyses of satellite (i.e. Landsat) or aerial photographs (e.g. Lepczyk et al. 2021), Digital Elevation Models (DEMs) have been long used for limited topographic analyses (e.g. Lepczyk et al. 2021). These span both Digital Terrain Models (DTMs; e.g. Melin et al. 2013), representing the elevation of a landscape as if stripped of any vertical surface features, and Digital Surface Models (DSMs; e.g. Lepczyk et al. 2021), which incorporate the elevation of the uppermost faces of features across a landscape surface. While useful for certain ecological questions at broad scales, a lot of structural complexity is unfortunately excluded by DEMs (e.g. Lepczyk et al. 2021). Therefore, for any research requiring fine structural details, terrestrial laser scanning and Structure-from-Motion photogrammetry are currently revolutionizing image processing into the truly 3D spatial realm (e.g. D’Urban Jackson et al. 2020; Lepczyk et al. 2021), and at a lower financial cost than other remote sensing data entails (e.g. D’Urban Jackson et al. 2020; Russo et al. 2023). The approach described in this paper utilizes the arguably more accessible (e.g. D’Urban Jackson et al. 2020) latter option, photogrammetry.

Photogrammetry (e.g. Granshaw 2015) refers to the process of 3D reconstruction from photographs, and has become popular for diverse applications, extending beyond academia (e.g. Pollefeys et al. 2004; D’Urban Jackson et al. 2020). The basic premise of photogrammetry is that overlapping images of a habitat or object will have identifiable features in common that can be matched, then stitched together to create a 3D scaffold; this scaffold can then be overlain with the original images to produce a textured model of the spatial scene. Photogrammetry has been used for mapping topographies in various research fields, including for reconstructions of anthropological sites (e.g. shipwrecks: Yamafune, Torres and Castro 2017; human graves: Badillo et al. 2020), for surveys of seafloor structure (e.g. Pollio 1968; Ciani et al. 1971; Hodúl et al 2018), and for the ecological and structural monitoring of different marine habitats (e.g. Roelfsema et al. 2022; Bayley et al. 2023; Yuval et al. 2023; Remmers et al. 2024; Paterson et al. 2024; AIMS LTMP 2025).

Importantly, only unobtrusive, user-friendly camera equipment, such as by GoPro, is needed to obtain the images for detailed photogrammetry models (e.g. Figueira et al. 2015; Guo et al. 2016), with such models boasting millimeter- to centimeter-level accuracy (e.g. Figueira et al. 2015; Guo et al. 2016; D’Urban Jackson et al. 2020). Studies in landscape and seascape ecology furthermore derive morphological measurements from photogrammetric models to estimate habitat complexity (e.g. Bayley and Mogg 2020; Lepczyk et al. 2021; D’Urban Jackson et al. 2020), thereby enabling comparisons between timepoints or locations with ecological interpretations (e.g. Yuval et al. 2023; Bayley and Mogg 2023). Three-dimensional metrics and analyses are still being explored for their conceptual value and will likely be developed further based on ecological questions (e.g. Vivancos, Closs, and Tentelier 2017; Kedron et al. 2019; Remmers et al. 2024; Lepczyk et al. 2021; Yuval et al. 2023). Several free, open-source platforms have been developed to enable photogrammetry (e.g. Colmap: Schönberger and Frahm 2016, Schönberger et al. 2016; Meshlab: Cignoni et al. 2008), further lowering the cost barriers associated with such methods, though a commercial software, Agisoft Metashape (Agisoft Developers 2025) seems more widely used for coral reef science. Accessible tools like these raise the tantalizing prospect of combining 3D geometric models of habitats with models of the sensory information available to animals as they move through 3D space.

Sensory ecology studies have recently begun addressing the combined challenges of tracking animals into reconstructed habitats, especially in the case of terrestrial insect research (e.g. Stürzl et al. 2015; Haalck et al. 2020; Haalck et al. 2023). For ants, Haalck et al. (2020) and Haalck et al. (2023) developed a method for automatically tracking 2D trajectories, across moderate distances, from the same footage used to reconstruct a map of the habitat. This automatic tracking method can handle target occlusion, inconsistent lighting, and poor contrast, however the animals must be filmed from a top-down perspective and at a constant distance from the camera (e.g. Haalck et al. 2020). Interestingly, Stürzl et al. (2015) reported a unique use of 3D habitat reconstruction in a study of 2D ant navigation, for which images from a color camera and laser scanner were combined in 3D to interactively model visual panoramas available to the ants at different locations along foraging routes. The same study recorded the 3D flight paths of ground-nesting wasps with a stereo camera system and registered these trajectories into a 3D reconstruction of the nest area, requiring visible calibration markers (Stürzl et al. 2015). Later, this was followed by a demonstration of how to quantify potentially available navigation information from 3D panoramas for flying insects (Murray and Zeil 2017).

Flying animals have been tracked in the field with several camera-based 3D tracking methods (e.g. Shelton, Jackson, and Hedrick 2014; de Margerie et al. 2015; Jackson et al. 2016; de Margerie, Pichot and Benhamou 2018), but typically by using fixed cameras and not including reconstruction of habitat structure, due to open backgrounds. Swimming animals have been tracked similarly (e.g. Vivancos, Closs, and Telier 2017). A particularly interesting approach from movement ecology tracked bats in 3D space using acoustic localization of their own calls and reconstructed the surrounding 3D vegetation structure with terrestrial laser scanning to investigate potential changes in flight pattern due to artificial light (Hermans et al. 2023).

Recently, habitat photogrammetry has been combined with 3D spatial tracking to study wild fish, both at the water surface (e.g. Choi et al. 2025) and underwater (e.g. Francisco, Nührenberg and Jordan 2020; Brady 2021). Francisco, Nührenberg and Jordan (2020) describe an approach that uses two or more cameras for tracking animals underwater in 3D and integrating these trajectories into 3D habitat reconstructions; unfortunately, the semi-automatic tracking cannot handle moving background objects, complex environments, animal occlusion, or highly variable lighting. Brady (2021) describes a method that uses two GoPro cameras for collecting separate photogrammetry and tracking footage, with a tracking procedure that consists of manual annotation of animal positions in both tracking videos. This approach is however reported as not suitable for long-range tracking, and a standardized object must remain within the video frames to also be annotated during tracking (Brady 2021).

To our knowledge, no studies have yet situated the 3D navigation movements of (underwater) animals in a detailed 3D habitat structure that incorporates relevant external sensory information. Our paper presents a method to address at least the first two parts of this challenge, for focal animals that have received extensive attention for visual perception but little attention for their movements in the wild.

Adult peacock mantis shrimp, *Odontodactylus scyllarus* (Stomatopoda: Odontodactylidae), are benthic predators of the Indo-pacific region (Ahyong 2001). They are 53–153 mm in body length, found between 1.2 - 30m depth, and live in burrows under the substrate (Ahyong 2001). Mantis shrimp are highly visual animals (e.g. Cronin et al. 2014; Cronin et al. 2022), capable of visual navigation (Patel and Cronin 2020a; 2020b; 2020c), and possess visual systems so complex (e.g. Marshall et al. 1996; Cowles et al. 2006; Marshall, Cronin, and Kleinlogel 2007; Chiou et al. 2008; Thoen et al. 2014; Daly et al. 2016; Daly et al. 2017; 2018; 2019; Chiou and Wang 2020; Wang and Marshall 2025) that an integrated model of visual perception does not yet exist for any mantis shrimp species. Moreover, the movements of *O. scyllarus* adults seem necessarily influenced by the structure of their habitat, since they walk and swim under, around, or through coral structures (personal observations). Although these animals both ambulate and swim, they usually travel on or just above the surface of the substrate regardless (personal observations). Potential influences of three-dimensional spatial complexity have not yet been accounted for in studies of mantis shrimp navigation, nor has mantis shrimp navigation been analyzed under wild conditions.

With an objective of describing 3D spatial use by *O. scyllarus* individuals through a wild reef habitat, the most important uncertainty we faced was that the cumulative distances or depth ranges traversed by these animals during excursions have not yet been described. We also did not know how frequently or for how long mantis shrimp individuals would become occluded during maneuvers around or through habitat structures. These two considerations rendered tracking via fixed-camera footage impractical. Due to the various other limitations of the video-tracking methods discussed above, existing approaches seemed ill-suited to our research organism and objectives.

We therefore decided to collect opportunistic tracking footage of mantis shrimp by following individuals with single GoPro cameras whilst SCUBA diving. To our knowledge, there is no tool currently available for automatically tracking animals in 3D from single-camera footage as dynamic as ours; the subject, background, and foreground all move and change, while the distance between camera and subject is ever inconsistent. We thus demonstrate a manual approach for precisely plotting the trajectories of mantis shrimp into a 3D reef map produced via photogrammetry.

This paper describes a straightforward method for reconstructing a 3D volumetric reef model (ca. 180 m^2^) and tracking the movement trajectories of peacock mantis shrimp from opportunistic field videos. Our approach is presented in four sections that span (1) collection of two types of GoPro camera footage whilst SCUBA diving, one for photogrammetry and the other for tracking animals, (2) photogrammetry of a measurable 3D reef habitat using popular commercial software, (3) example morphological measurements of habitat complexity from the finalized 3D model, and (4) manual plotting and measurement of animal trajectories on the surface of the 3D model.

Tracking methods have been acknowledged to bias larger animals in movement ecology (e.g. Kays et al. 2015), and seem to bias smaller animals in sensory ecology, though this disparity will gradually shrink as technology continues to advance (e.g. Kays et al. 2015). At present, invertebrates that live underwater and/or in structurally complex habitats seem among the least likely to be tracked in the field. To encourage contextualized 3D movement studies of organisms that have received little or no attention in the wild, researchers should have access to simple methods that rely on minimal assumptions about movement patterns, require minimal field equipment or personnel, and do not require much extra time spent at the field site. Our approach satisfies all of these concerns, so we expect that it could be adapted for studies of diverse animals, whether in aquatic or terrestrial habitats.

## Data Collection

Plotting animal trajectories in 3D requires two types of footage, one for creating a 3D photogrammetric model of the habitat, and one that records the translational movements of the animals. This paper refers to videos that follow moving animals as *Animal Footage* and to videos (or photos) that capture the detailed habitat for photogrammetry as *Habitat Footage*. In our case, both types of footage were obtained in video (mp4) format, using GoPro Hero 11 cameras while SCUBA diving in a team of two.

Our fieldwork was conducted 7^th^ to 24^th^ November 2024 at the House Reef of Kuredhdhoo (Kuredu Island Resort), Lhaviyani Atoll, Maldives, under permit number NRP2024/101 (Ministry of Fisheries and Ocean Resources, Republic of Maldives).

House Reef is a fully submerged patchy reef that faces the inside of the atoll ring and experiences a prevailing current in the SW-NE direction. We found four peacock mantis shrimp (*Odontodactylus scyllarus*) individuals living in the same ca. 180 m^2^ coral reef patch that was located < 100 m east of the end of Kuredu Island Jetty. This sloping reef patch spanned ca. 8 - 13 m depth, was largely characterized by coral rubble, and was bordered on all sides by coral sand.

Our two researchers spent 18 days (31 dives) of active field time at Kuredhdhoo. Of these, 17 days (29 dives) were dedicated to obtaining *Animal Footage*, and 1 day (2 dives) was dedicated to collecting *Habitat Footage*.

### Animal Footage

To collect *Animal Footage* of translationally moving animals, we used two GoPro Hero 11 cameras, each in a waterproof housing fastened to a selfie stick (49.4 cm or 94.5 cm long, including the GoPro mount) such that camera lenses pointed away from users. The GoPro Hero 11s were set to 4K resolution, 24 fps, with a wide-field view, and data was written to 128GB SanDisk Extreme microSD cards, to be copied onto external hard drives nightly.

This extendable camera configuration allowed both SCUBA divers to maintain a greater distance from the mantis shrimp we were following. Over 17 days dedicated to finding animals and collecting *Animal Footage*, we conducted up to two buddy dives per day, to 11.0 ± 1.7 m max depth, resulting in 29 total dives that lasted 93.3 ± 16.6 minutes. We found four peacock mantis shrimp living in the same ca. 180 m^2^ of sloping reef, which spanned about 8 - 13 m depth.

Each dive began with a preliminary visual survey to see whether any previously identified animals were out of their burrows already. If not, each diver chose one mantis shrimp burrow to watch from ≥ 2m distance, keeping a low profile. When a mantis shrimp left its burrow, the diver filmed it with the GoPro on a stick until either the animal returned to its burrow, the diver lost sight of it and could not find it again, or the divers had to end their dive due to low air. Animals were followed and filmed from behind or from the side, depending on the water current, the reef topography, and the overall slope of the reef. While following animals, we sought to minimize potential influence of our presence on the animals by keeping our bodies low (≤ 30° above an individual, relative to the substrate), and maintaining a distance of ≥ 1 m between ourselves and the mantis shrimp, adjusting based on perceived boldness/shyness or signs of sudden alertness.

If a mantis shrimp was found outside of its burrow, it was also filmed until either it returned to the burrow, the diver lost it, or the dive had to end. As more data was collected, the divers prioritized waiting longer for certain individuals, based on how much *Animal Footage* had already been collected for each animal. In total, we collected approximately 20 hours (381.21 GB) of *Animal Footage*, and this comprised 121 separate excursions, though unequally represented by the four individual animals.

### Habitat Footage

To create a scalable 3D model of a habitat, overlapping videos or photos must be collected such that the entire area of interest is captured in the *Habitat Footage*. Additionally, markers of a known size must be placed within the habitat before it is recorded. Our approach to collecting *Habitat Footage* was inspired by Roelfsema et al. (2022, section 2.4.1.) and van den Berg et al. (2022). The most notable differences include that our method uses two GoPro cameras instead of three, and our original *Habitat Footage* was collected in video format, rather than as photographs.

The first step is to construct a *Photogrammetry Rig*, which functions to keep the cameras at a constant orientation relative to each other. Our *Photogrammetry Rig* (Fig. 1 top right) was constructed from an aluminum tube (60.4 cm x 5.0 cm x 5.0 cm) and two GoPro Hero11 cameras in waterproof housings. The GoPro housings were attached to the tube at 3.4 cm from either end, centered on the midline of the width, and angled at 71° and 101° such that their fields of view overlapped (≥ 60% overlap, at a distance of 1.5 - 2m above the reef). Both cameras recorded videos at 5312 x 2988 resolution (5.3K) and 60 fps, in wide field.

**Figure 1:**
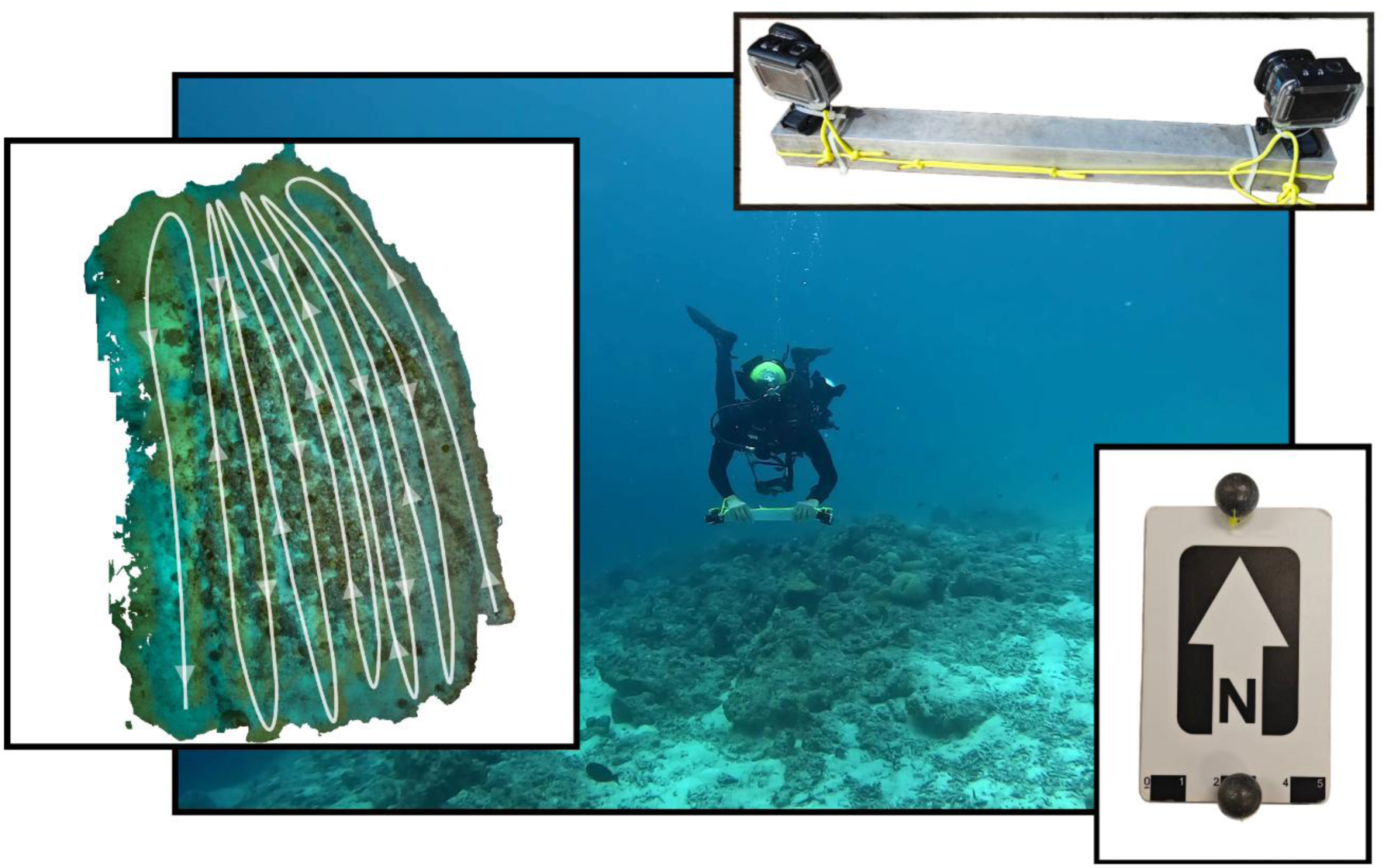
Swim-over filming and the equipment used to collect footage for 3D photogrammetric reconstructions of the reef habitat. (Left) Top view of the approximate swim-over pattern conducted while filming the reef, with arrow heads indicating the diver’s heading; re-visitations to large, shadowy objects occurred after this pattern was completed. The Photogrammetry Rig (top right) was constructed from a length of square aluminum tube (60.4 cm x 5.0 cm x 5.0 cm) and two GoPro 11 housings with overlapping fields of view. Yellow paracord was looped through the tube to provide an attachment point for the diver and to secure the cameras in the event that attachments failed. Five Photogrammetry Markers (bottom right; van den Berg et al. 2022) were distributed in the reef scene prior to data collection. Each Marker displayed an arrow to indicate north and a five-centimeter ruler at one short edge. Markers were kept negatively buoyant with one fishing weight fastened at either short edge. The back panel shows deployment of the photogrammetry rig during the swim-over sequence. One diver (M.S.) swam at neutral buoyancy with consistent speed, approximately 1.5 - 2 m above the reef, pointing the camera lenses downwards. Another diver (M.D.) served as an intermittent visual guide, just off of the bottom edge of the swim-over map (left). This photograph has been enhanced for clarity.

Our reef patch of interest covered an area of approximately 180 m^2^ and was situated on a slope, spanning about 8 - 13 m depth. Before collecting *Habitat Footage*, we placed five black and white *Photogrammetry Markers* (Fig. 1 bottom right; courtesy Cedric van den Berg; van den Berg et al. 2022; Roelfsema et al. 2022) approximately equidistant from each other, near the four corners and center of the reef habitat. Each marker featured a 5cm ruler and an arrow, which was pointed towards magnetic North by consulting a dive compass. Fishing weights, fastened to both short ends of each *Photogrammetry Marker*, kept the markers negatively buoyant and stationary.

Once the *Photogrammetry Markers* had been placed, a swim-over sequence was conducted, similar to Roelfsema et al. (2022, Fig. 10D). The diver holding the *Photogrammetry Rig* started recording with both cameras, then began to steadily swim about 1.5 - 2 m above the reef, holding the aluminum bar such that the lenses of both cameras were oriented downwards (Fig. 1 back). The planned swimming pattern (Fig. 1 left) proceeded back and forth across the reef in approximately parallel lines separated by around 1.5 - 2 m. The second diver hovered ca. 3 m away from one edge of the reef patch, providing a visual target to keep the swim-over pattern on track. Each time the recording diver reached the near end of the reef and turned away, the visual-target diver moved approximately 1.5 - 2 m to the side for the next approach. Once the entire area had been filmed, the recording diver revisited any large, particularly complex, and/or shadowy structures present on the reef, filming them from as many angles as possible. This reduced the likelihood that the finished model would include any gaps.

Between both cameras, we collected approximately 2 hours total (51.2 GB) of *Habitat Footage* in .mp4 format, during ca. 3 hours of diving (1 day, 2 dives). Images were then extracted from the footage before proceeding with photogrammetric reconstruction.

## Photogrammetric Reconstruction

For photogrammetric reconstruction of the reef habitat, we used the commercial software Metashape Professional (version 2.2.2, Agisoft, St. Petersburg, Russia), under an Educational License. Since our Photogrammetry Footage had been collected in video format, images first had to be extracted for loading into the Metashape software. Images were extracted at 1 fps using FFmpeg (FFmpeg Developers 2025) with the line of code below:

ffmpeg -I /pathandnameof/videofile.mp4 -r 1 /savedirectoryand/filename_%04d.png

Extraction resulted in 6683 total images (185.27 GB) in .png format, whereof the images from the parallel swim-over pattern demonstrated an overlap of ≥ 60% between paired images. All images were added to a new Metashape project for processing. Neither the videos nor the images needed to be synchronized for any part of this process.

The basic workflow in Metashape calculates tie points in common between overlapping images, creates a three-dimensional cloud of these common points, builds a solid 3D model connecting the points, and finally overlays the colors and textures from the originally uploaded images (Agisoft Developers 2025) (Fig. 2).

**Figure 2:**
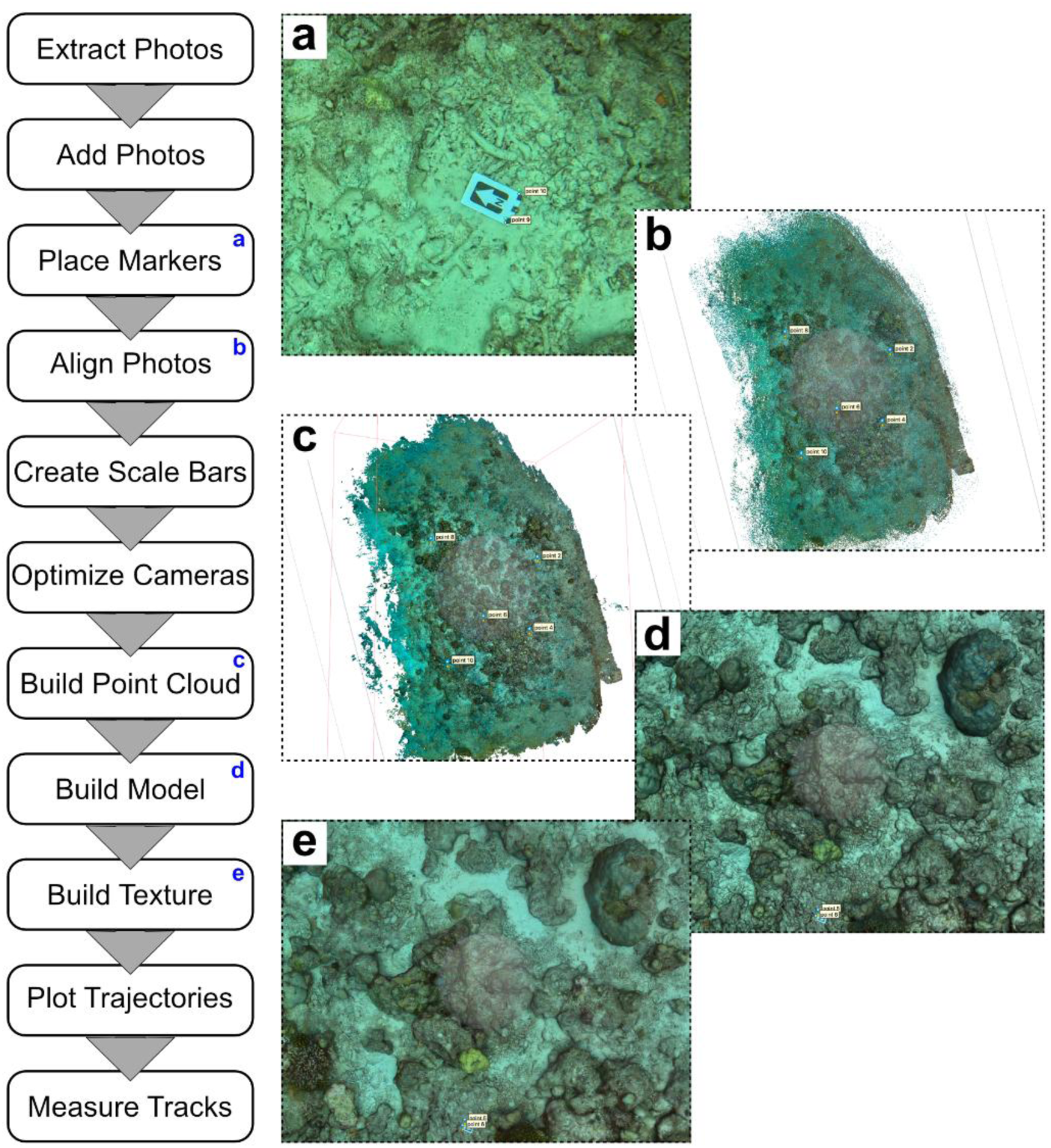
Workflow for creating a 3D reef model. The first, *Extract Photos* step is achieved using Ffmpeg, while all of the following steps are completed in Agisoft Metashape Professional. The left column depicts the steps of the workflow, and the right column illustrates five selected steps. Blue letters in the left column corresponding to the letters in the illustrations on the right. An optional and subjective cleanup step may be included between *Build Texture* and *Plot Trajectories* (see text).

Immediately after adding images into the Metashape project, “marker” points must be placed at either end of each ruler on each Photogrammetry Marker in all images that clearly feature a Photogrammetry Marker (Fig. 2a, *Place Markers*). The same pair of point identifiers must be used for each given Photogrammetry Marker in all images where that particular Marker is annotated. We used five Photogrammetry Markers, so ten points were repeatedly placed. Once the Markers are annotated, the images are aligned (Fig. 2b, *Align Photos*), which creates a sparse cloud of tie points based on estimated camera positions. Then, scale bars must be created for each pair of points, a step that is essential when building a model that is scalable and measurable. Following the creation of scale bars, the *Optimize Cameras* step triangulates tie point coordinates based on any available measurement information, including camera positions, placed marker coordinates, and scale bar distances. Following this order of steps is therefore important for maximizing geometric accuracy in the final model (Agisoft Developers 2025).

After *Optimize Cameras*, the next step is *Build Point Cloud* (Fig. 2c), which creates a dense cloud of triangulated tie points. During this step, “Depth Maps” are calculated and combined using orientation information from overlapping images. Once the point cloud has finished processing, noise may be deleted anywhere in the 3D cloud by manually selecting groups of unwanted points with the Selection Tools and deleting them. However, if the point cloud itself will not be used as the Source Data during the next *Build Model* step, deleting noise from the point cloud will not save any processing time.

The *Build Model* step (Fig. 2d) creates a solid polygonal model. We used “Depth Maps” for our Source Data because this is recommended for “Arbitrary” surface types (Agisoft Developers 2025); note that these Depth Maps will still include any tie points that were deleted from the point cloud. After *Build Model*, the final step is *Build Texture* (Fig. 2e), which applies color textures to the model based on the original images, or other selected Source Data options. In case lighting was not consistent during data collection, a lengthy *Color Calibration* step can be included if required, but this must be done before the *Build Texture* step (Agisoft Developers 2025). Any remaining noise in the model that consists of isolated, floating fragments can be quickly removed from the final model by using the *Clean Model* tool (“Tools” > “Model” > *Clean Model*) and setting the slider to 99%. Holes may then be filled using the *Close Holes* tool, as explained later.

Throughout the Metashape workflow, the user is asked to select from certain parameters before processing commences. Different options for Accuracy/Quality settings at each step of the workflow determine whether the software uses the original images versus downscaled or upscaled versions of the images (Agisoft Developers 2025). Furthermore, during the *Build Point Cloud* and *Build Model* steps, the Depth Filtering parameter determines the rigorousness with which outlying points are removed (Agisoft Developers 2025). Although the Metashape manual does not recommend the default “aggressive” depth filtering for highly textured scenes or scenes with fine details (Agisoft Developers 2025; Over et al. 2021), “aggressive” was the appropriate setting for our purposes; this resulted in the least noise while retaining a high level of detail.

Our reconstruction of the reef patch was made using “high” accuracy/quality settings at each step of the workflow, which resulted in a reconstruction that was already detailed and suitable for plotting animal trajectories. With the parameters we used (Table 1), the workflow took about eight days of processing time on a Mac Mini 2020 with Apple M1 chip, Sequoia 15.7.2 operating system, and 16 GB memory.

**Table 1:**
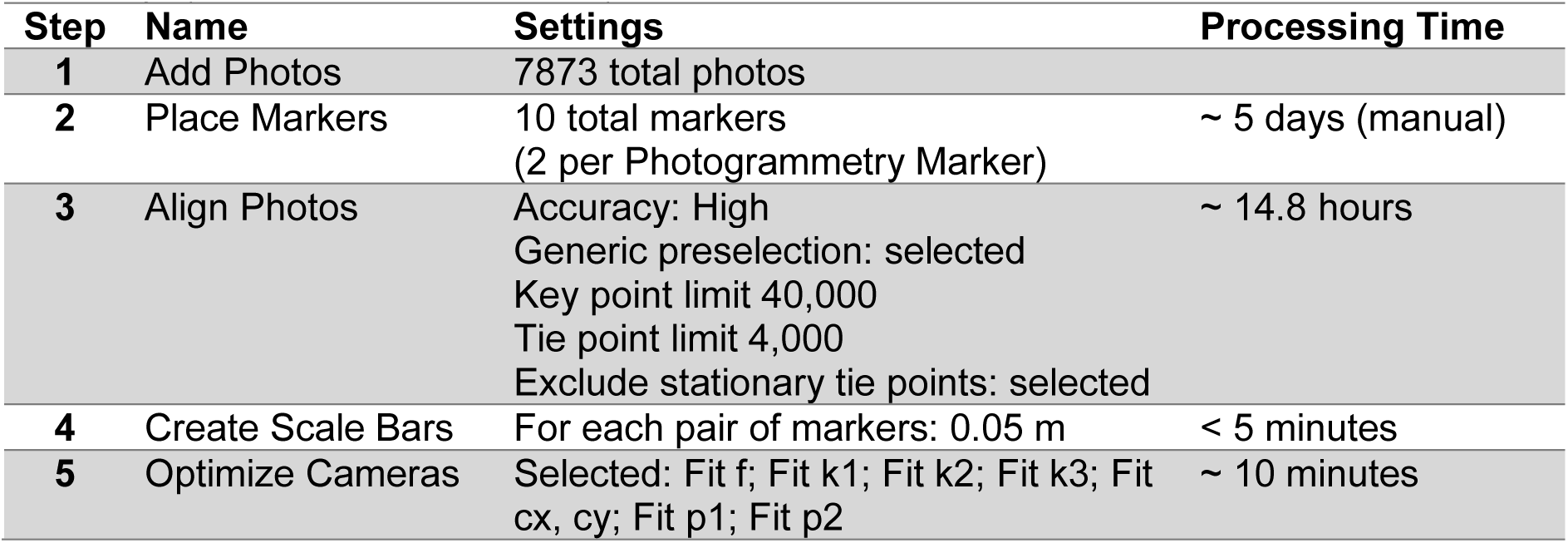
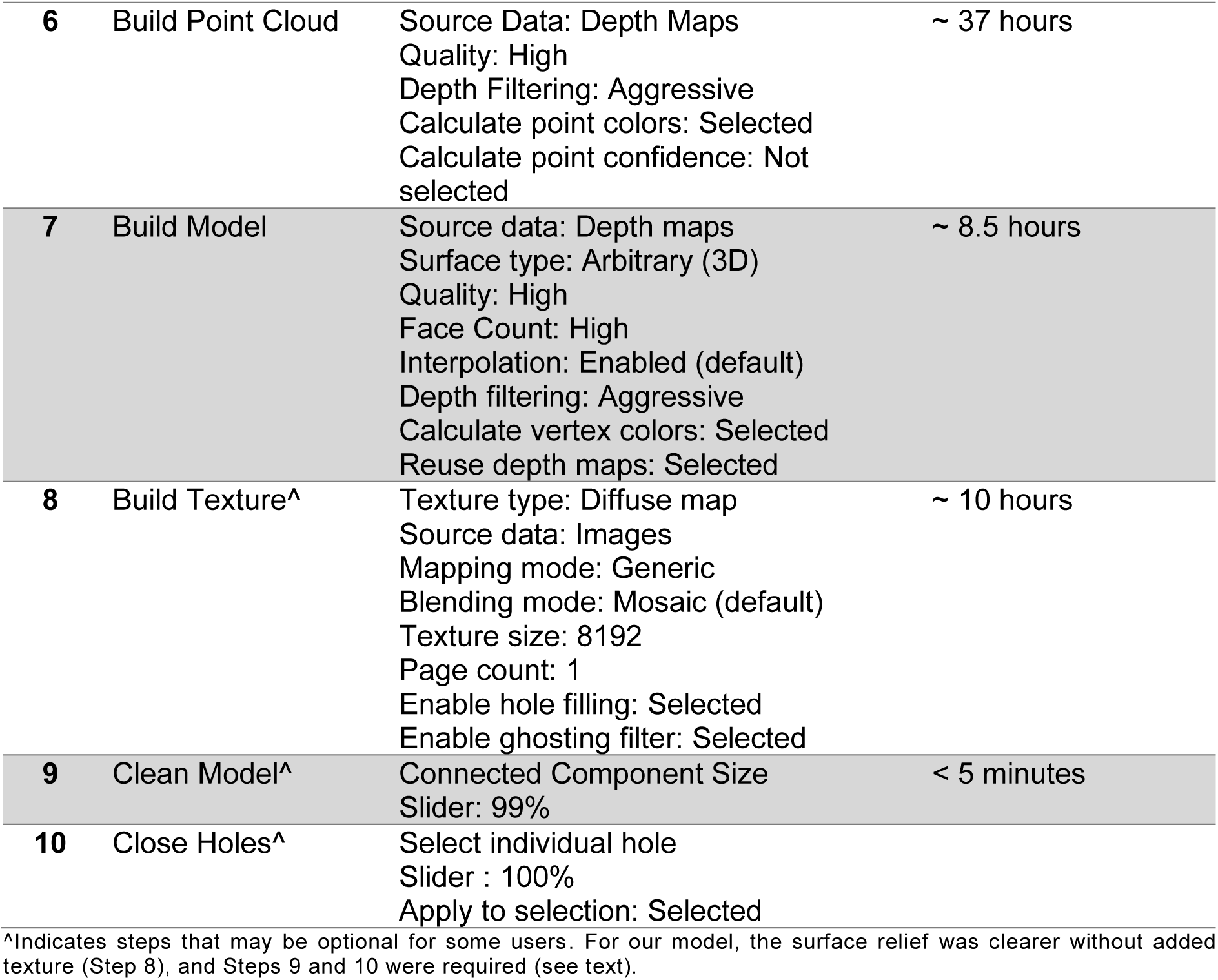
Photogrammetry parameters for the final reef model. Settings used for each step of the photogrammetry workflow in Agisoft Metashape Professional, version 2.2.2 (2005). The associated processing times are included for a Mac Mini 2020 with Apple M1 chip, Sequoia 15.7.2 operating system, and 16 GB memory.

| Step | Name | Settings | Processing Time |
| --- | --- | --- | --- |
| 1 | Add Photos | 7873 total photos |  |
| 2 | Place Markers | 10 total markers<br>(2 per Photogrammetry Marker) | ~ 5 days (manual) |
| 3 | Align Photos | Accuracy: High<br>Generic preselection: selected<br>Key point limit 40,000<br>Tie point limit 4,000<br>Exclude stationary tie points: selected | ~ 14.8 hours |
| 4 | Create Scale Bars | For each pair of markers: 0.05 m | < 5 minutes |
| 5 | Optimize Cameras | Selected: Fit f; Fit k1; Fit k2; Fit k3; Fit cx, cy; Fit p1; Fit p2 | ~ 10 minutes |
| 6 | Build Point Cloud | Source Data: Depth Maps<br>Quality: High<br>Depth Filtering: Aggressive<br>Calculate point colors: Selected<br>Calculate point confidence: Not selected | ~ 37 hours |
| 7 | Build Model | Source data: Depth maps<br>Surface type: Arbitrary (3D)<br>Quality: High<br>Face Count: High<br>Interpolation: Enabled (default)<br>Depth filtering: Aggressive<br>Calculate vertex colors: Selected<br>Reuse depth maps: Selected | ~ 8.5 hours |
| 8 | Build Texture<sup>^</sup> | Texture type: Diffuse map<br>Source data: Images<br>Mapping mode: Generic<br>Blending mode: Mosaic (default)<br>Texture size: 8192<br>Page count: 1<br>Enable hole filling: Selected<br>Enable ghosting filter: Selected | ~ 10 hours |
| 9 | Clean Model<sup>^</sup> | Connected Component Size<br>Slider: 99% | < 5 minutes |
| 10 | Close Holes<sup>^</sup> | Select individual hole<br>Slider : 100%<br>Apply to selection: Selected |  |

<sup>^</sup>Indicates steps that may be optional for some users. For our model, the surface relief was clearer without added texture (Step 8), and Steps 9 and 10 were required (see text).

The resulting model with texture is shown in Figure 3 as an orthomosaic projection. Our final model included nine noticeable holes, which we filled using the *Close Holes* tool (Fig. 4). We found it best to select individual holes and fill them each to 100%, the whole process of which took less than a minute per hole. When we tried to close holes for the entire model at once, any indentations in the model were also closed such that there were flat surfaces where there should not have been, probably as a function of the automatically generated local coordinate system of our model. It is worth noting that the *Close Holes* tool results in a solid color filling (Fig. 4c), with no obvious texture to match the surrounding surface, so it is still best to avoid large holes as much as possible if texture is important.

**Figure 3:**
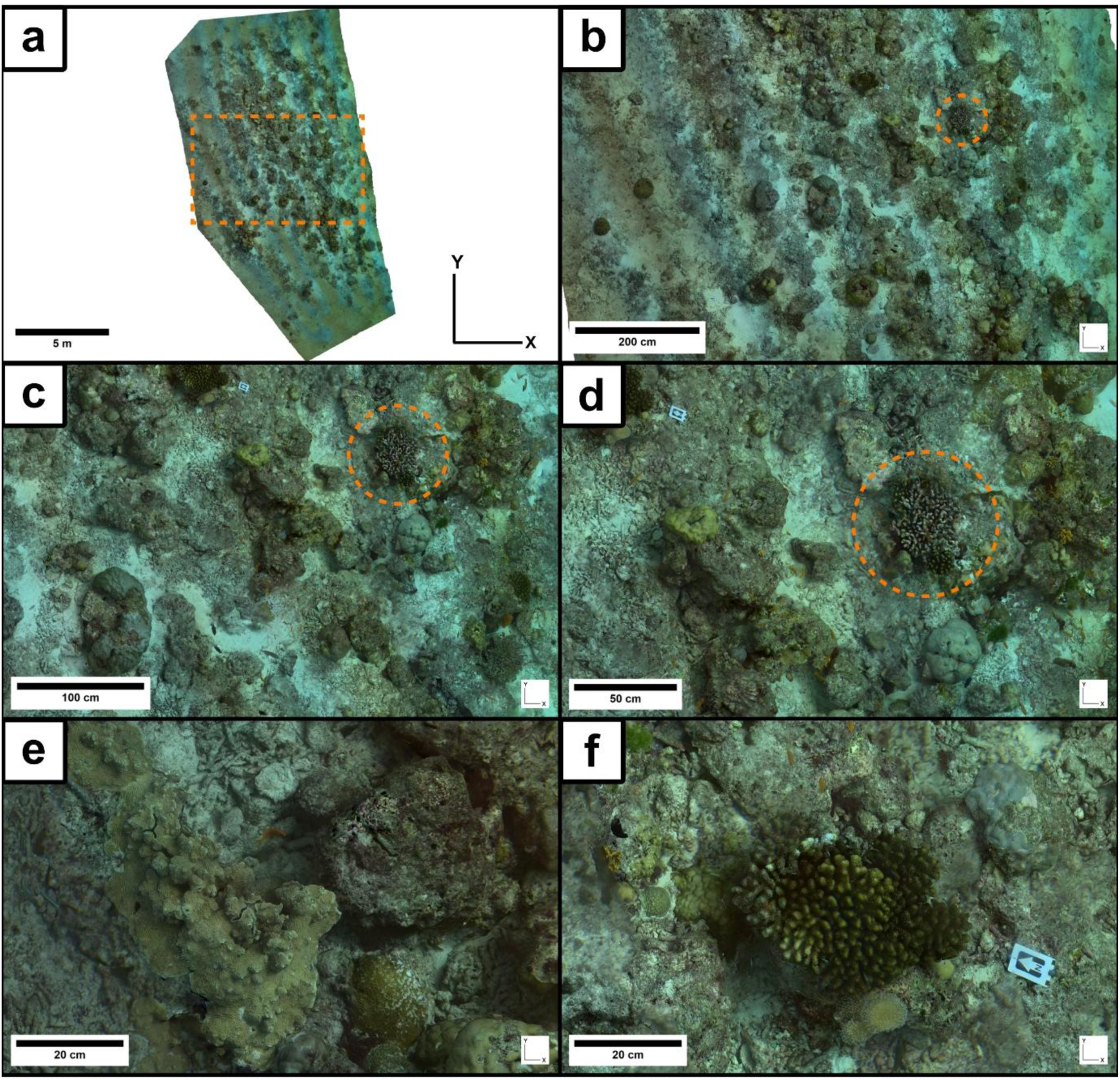
Final model with texture, shown as a 2D orthomosaic map in the xy plane. (a) The whole 2D map of the reef, with a dotted orange box delineating the area shown in panel (b). An enlarged xy-compass is included for orientation. Panels (b-d) show increasing magnifications within the same region, with a dotted orange circle surrounding an example coral head that is visible in each panel. Panels (e) and (f) show different example regions at the same high magnification to each other. The region shown in (e) is external to the region shown in (b), while the region shown in (f) is present in (b-d), identifiable by the photogrammetry marker. Note that the scalebars show different distances as magnification increases.

**Figure 4:**
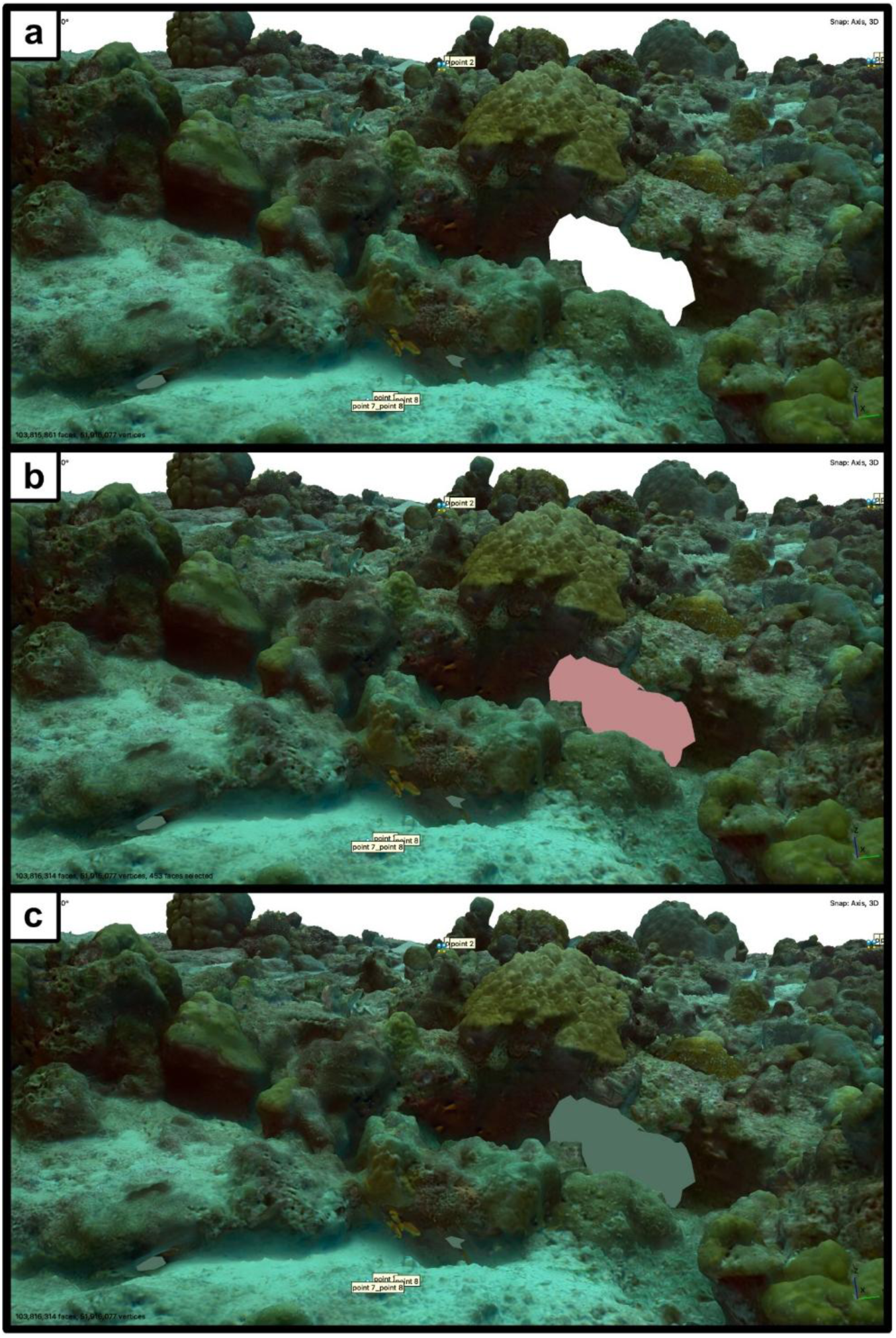
An example hole before and after closing. (a) The largest hole in the reef model. Holes are faceless gaps in the model surface, identifiable by the white showing through from the empty space around the model. (b) Immediately after using the *Close Holes* tool to fill the largest hole. The filling is automatically highlighted in a rose color, indicating it is selected. In the bottom left corner of the screen is displayed that this selection comprises 453 faces. (c) The largest hole after closing, showing the uniform green color of the filled area when not selected. Notice the two smaller holes that have also been closed, in the same uniform green color. Some fish have been incorporated into the texture projected onto the model because they did not move while their region was being filmed.

Users may encounter surface noise in their final models, which may be problematic for tracking animals later, if the animals traversed those areas of the habitat in which the surface noise occurs. Removing surface noise and closing the resulting holes can restore a relief to the model that is better representative of the surface in reality. If a habitat model will also be used to derive measurements for describing habitat complexity, substantial noise must be removed/corrected to improve measurement accuracy.

In our case, the underside of the final model showed several long, artificial tubes/tunnels leading away from small would-be holes. Since these artificial tunnels would influence our measurements of 3D surface area or volume, we removed them manually, by repeatedly selecting parts of a given tunnel with the Selection tool and then deleting the selected faces. Though these tunnels originally had closed ends, removing them necessarily opened small holes in the model surface, which then had to be patched using the *Close Hole* tool.

Before removing noise and closing holes, our model consisted of 115,145,953 faces and 57,606,987 vertices. The cleaned and patched model consisted of 103,816,314 faces and 51,916,077 vertices. The 3D surface area of our finished model was 272.65 m^2^, and the fitted 2D surface area was 184.07m^2^.

Our model provided more obvious relief shadowing without texture than with texture, so removing texture allowed for more intuitive visual judgments of relative distance, for the purpose of manually plotting trajectories. Texture can be removed from a model in less than two minutes by right clicking on the model in the Workspace pane and selecting “Remove” > “Remove Texture”. We recommend duplicating the model first, by right clicking and selecting “Duplicate”.

Finally, in order to move on to plotting trajectories in the model or creating orthomosaic projections, the coordinate system must be selected and set in the Reference Settings. We set our system to “Local Coordinates (m),” since we did not have the means to impose an external coordinate system on our model.

## Measuring the 3D Model

There are many ways that ecologists quantify surface topography (e.g. Bayley and Mogg 2020), including rugosity (e.g. Knudby and LeDrew 2007) and fractal dimension (e.g. Bradbury and Reichelt 1983; Yuval et al. 2023), as well as Surface Area/Volume (e.g. Yuval et al. 2023). Such measurements allow ecologists to objectively compare habitats with similar structural elements and even draw conclusions about the relative health of their ecosystems. We offer two example measurements of complexity from our model reef.

Rugosity is the measurement of structural complexity used most commonly for coral reefs (Knudby and LeDrew 2007). This measurement is traditionally based on linear transects, calculated by draping a length of a chain or line across a transect and dividing the total length used by the straight-line distance measured between the start and end points of the transect (Knudby and LeDrew 2007). Rugosity thus usually depends on the method used as well as substrate type and scale (Knudby and LeDrew 2007); relatedly, fractal dimension describes how rugosity changes across scale (Knudby and LeDrew 2007). One way to calculate rugosity from a 3D habitat model is to divide its 3D surface area by the 2D surface area of the leveled plane on which it sits (e.g. Roelfsema et al. 2022). To calculate the rugosity of our 3D reef model, we measured the 3D surface area of the model via *Tools ◊ Model ◊ Measure Area and Volume*, then measured the 2D surface area (“Area, fitted 2D”) by outlining the model with the *Draw Polygon* tool, right-clicking on the shape, and selecting *Measure.* With a 3D surface area of 272.65 m^2^ and a 2D surface area of 184.07 m^2^, our model’s rugosity is 1.48.

Surface Area/Volume is another way to measure surface complexity, where higher numbers correspond to more complex structures (Yuval et al. 2023). In order to calculate Surface Area/Volume, we first created a duplicate of the model, closed the duplicate model to 100% using the *Close Holes* tool, and then measured the 3D surface area and volume via *Tools ◊ Model ◊ Measure Area and Volume*. With a surface area of 461.17 m^2^ and a volume of 55.72 m^3^, our closed model’s Surface Area/Volume is 8.28 m^-1^.

## Plotting and Measuring Trajectories

To our knowledge, no existing tools are able to automatically track animals through extensive video footage in which all regions, dimensions, and aspects of the footage are in flux. We therefore propose manual plotting of trajectories into 3D models as a way to track animals through complex habitats.

Once a 3D model of the habitat has been completed, it is possible to plot the trajectories of animals on the surface of the model, within Metashape Professional (version 2.2.2, Agisoft, St. Petersburg, Russia). This is done by alternately watching *Animal Footage* and plotting points into the 3D model that correspond to the location of the animal in the video. The changing structural features of the habitat that are present in the *Animal Footage* should also be present in the model, and recognizing these key features while manipulating the model perspective makes it possible to determine an animal’s precise location and position throughout the video sequences.

We plotted trajectories using the *Draw Polyline* tool (Agisoft Developers 2025). For longer trajectories and/or *Animal Footage* with variable viewing angles, it is necessary to zoom in/out and rotate the 3D model so that features can be viewed from different perspectives. Unfortunately, the *Draw Polyline* tool does not allow the user to manipulate the 3D model while plotting consecutive points of a polyline, but it does allow the user to add points (“vertices”) along an existing line and also to move vertices that already exist. It is therefore possible to plot long and complex trajectories by first plotting just three or four initial points (Fig. 5a), one at the approximate start of the trajectory, one or two somewhere in the middle, and one near the end of the trajectory. Once these guide points are plotted, completing a rough, initial polyline, the trajectory can be edited into all its complexity by adding vertices and repositioning them as needed (Fig. 5b). This approach allowed us to work around the limitations of trying to consecutively plot long (20+ m) polylines in one go, since we were able to manipulate the model in between adding or dragging vertices. Figure 5b and d show the same intermediate example step in trajectory plotting, just from different angles.

**Figure 5:**
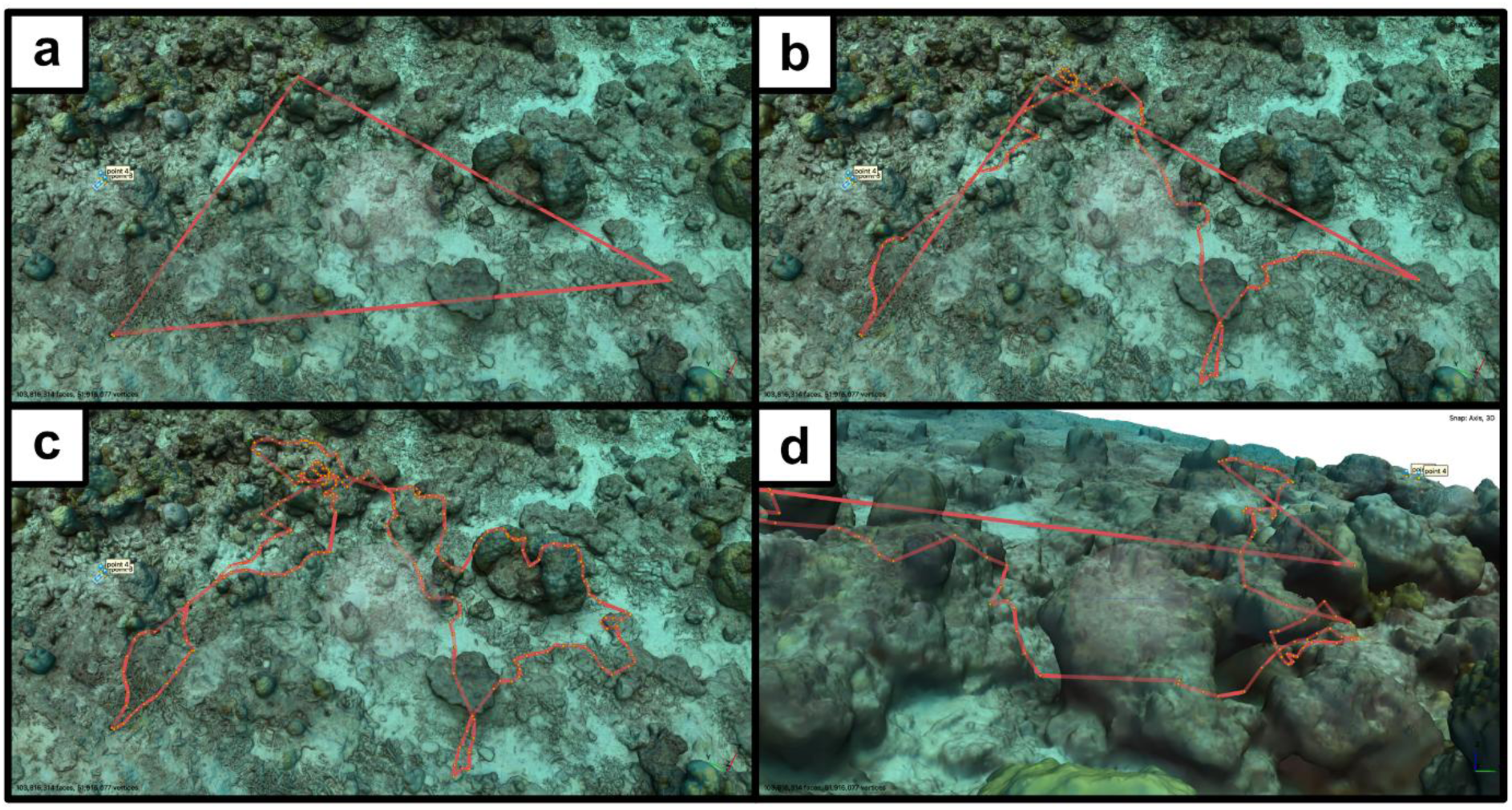
Example trajectory of a mantis shrimp plotted into the 3D reef model. Panels a-c show the general stages of the plotting process, all from the same viewing angle and magnification. (a) shows the rough initial polyline, (b) shows the trajectory about halfway through plotting, as vertices are being added to the polyline, and (c) shows the completed trajectory, which measures 20.01 m in 3D and 17.67 m in 2D. (d) Another angle of view on the model with the trajectory at the same stage as in (b) during plotting: The model is oriented to match the angle of view shown by a frame of video in the *Animal Footage*.

Once finished, the entire polyline trajectory (Fig. 5c) is then immediately measurable in both 2D and 3D by right-clicking on the trajectory and selecting *Measure*. If smaller portions of a larger trajectory are of interest, the *Ruler* tool can be used to trace that portion of the trajectory, returning an immediate value. Multiple trajectories can be grouped and color-coded for individual animals. Once trajectories are completed, it is possible to export these as “shapes” with .shp, .dxf, .gpkg, or .txt formats for further analyses in external software. For models that use an external coordinate system, .kml, .kmz, and .geojson are also available export formats for shapes.

The nature of our *Animal Footage* meant that animals were not continually visible. Whenever a mantis shrimp became occluded, for example when it disappeared under or behind a coral head, vertices were added in an approximately straight line connecting the last visible location to the position of the animal once it became visible in the Tracking Footage again. Furthermore, whenever an animal left the substrate surface to swim between two locations of different depth, a vertex was added at the position of the animal’s last contact with the substrate, and another vertex was added to the location at which it re-established contact with the substrate.

## Discussion

The method we have described includes three general parts: filming the navigation paths of wild animals by following them at a distance, reconstructing a three-dimensional (3D) spatial habitat model from separate footage, and manually plotting the 3D trajectories of animals into the digital reef model. Our approach includes (1) collection of GoPro camera footage whilst SCUBA diving, one kind of footage for photogrammetry and the other for tracking animals, (2) photogrammetry of a measurable 3D reef habitat using Agisoft Metashape Professional (version 2.2.2, Agisoft, St. Petersburg, Russia), (3) example morphological measurements of habitat complexity from the finalized 3D model, and (4) manual plotting of animal trajectories, onto the 3D model and measurement of the trajectory lengths as 3D and 2D distances. Our example animal is the peacock mantis shrimp (*Odontodactylus scyllarus*), a marine invertebrate, but this methodology will be adaptable for other aquatic and terrestrial study systems because the main factors that inspired our approach include: the common limitations of existing tracking methods (e.g. Vivancos, Closs, and Telier 2017; Francisco, Nührenberg and Jordan 2020; Haalck et al. 2020; Brady 2021), the understudied status of our focal species in its wild context, and the logistical challenges of fieldwork (i.e. financial expense, field time, equipment capacity, variable weather conditions, and 3D habitat structure).

First, filming animals with small cameras on sticks allows researchers to flexibly adjust the distances and angles of themselves and their cameras relative to focal individuals or field obstacles, without having to know the range or route of the animal in advance. To our knowledge, this kind of footage is not useable with currently available 3D tracking software (e.g. Vivancos, Closs, and Telier 2017; Francisco, Nührenberg and Jordan 2020; Brady 2021), but filming in this way would allow researchers to reduce the likelihood of their presence influencing animal behavior, while also enabling them to maneuver around obstacles and occlusions arising along 3D animal trajectories. In our case, an unanticipated benefit of obtaining *Animal Footage* in this way was that we were able to record and document rarely observed and previously undescribed behaviors for our species (not yet published).

Next, the photogrammetry equipment we used was influenced by airline requirements and baggage costs. We had to use a shorter *Photogrammetry Rig* than Roelfsema et al. (2022), since it had to fit in a suitcase, so we mounted just two GoPro cameras rather than three. This also reduced the amount of data collected for *Habitat Footage*.

The format of *Habitat Footage* which we recommend was influenced by limited field time and unpredictable field conditions. Fieldwork is often unpredictable, and in our case data collection took place during the transition out of monsoon season in the Maldives, so changes in weather and their effects on underwater currents and visibility were unforeseeable. Our decision to collect videos, rather than images, for *Habitat Footage* reduced the likelihood that accidental fluctuations in swimming speed or depth would be problematic for data processing later. We therefore did not need to calculate and practice the required swimming speed that would result in enough overlap between timelapse images. Instead, it was possible for us to return from the field and extract images at different frame counts per second (fps) to determine which fps resulted in images with an appropriate amount of overlap. Collecting *Habitat Footage* in this format will minimize the total time devoted to collecting such footage, as well. Researchers whose computers are more limited in processing power may need to determine the minimal number of images or the minimal percentage of overlap that would produce a viable photogrammetric model under their given field conditions. We would therefore advise anyone who has only one or few chance(s) to collect *Habitat Footage* from a field site that they do so in video format.

The photogrammetry workflow (Fig. 2) we provided for creating a 3D reef model in Agisoft Metashape Professional version 2.2.2 (2025) is straightforward. The Metashape User Manual (Agisoft Developers 2025) is certainly useful, though it spans 240 pages and is written in a way meant to cover all use-cases, so interpreting and trialing the software for our use-case was time-consuming and repetitive. We hope that including the details of our final procedure (Table 1) has made the photogrammetry software more accessible and encouraging to researchers interested in studying animal movements and interactions in 3D space.

From our final model, we chose to include two morphological measurements of structural complexity because reporting such measurements might aid comparability as movement ecology and sensory ecology begin to contextualize animal movements into wild, 3D habitats. By using similar data collection procedures and processing parameters, researchers could compare how different structural complexities at different times, under different conditions, or at different field sites influence movement patterns within or between different species. Since coral reefs are generally diverse and complex, it should be noted that single measurements will not be comprehensively useful for comparing structure across scales and habitats (Knudby and LeDrew 2007). Given the historically 2D or 2.5D perspective of landscape ecology (e.g. Lepczyk et al. 2021), tools and concepts for 3D morphometrics are still developing (e.g. Vivancos, Closs, and Tentelier 2017; Kedron et al. 2019; Lepczyk et al. 2021; Yuval et al. 2023), so perspectives from movement ecology (e.g. Nathan et al. 2008) and sensory ecology (e.g. Dangles et al. 2009; Willemart 2023) could be useful in refining them or expanding their applicability. We encourage interdisciplinary communications and collaborations.

Finally, we described a method of tracking animal trajectories that is heavily manual. Ideally, future automatic/semi-automatic tracking software will be developed that can handle the extremely dynamic format of *Animal Footage* that we presented, but for now manually plotting trajectories into 3D models in Agisoft Metashape Professional version 2.2.2 (2025) offers a precise way to advance research for understudied animals. The most important limitation of plotting trajectories in this way is probably that it is not possible to plot points in the empty space of the model in this software, only on the model surface. Users will therefore not be able to track animals using our approach if the animals swim or fly considerable distances above the substrate. Mantis shrimp do swim but usually very close to the substrate, so it is still possible for us to plot their trajectories on the model surface in Metashape as a close approximation. For those researching benthopelagic animals, the 3D tracking methods developed by Francisco, Nührenberg and Jordan (2020) or Brady (2021) may be of interest, assuming research questions fall within the software limitations.

On the other hand, for those interested in benthic or terrestrial organisms known to move very slowly (e.g. centimeters to meters per year), the creation of multiple photogrammetric reconstructions, from data collected at disparate timepoints may, on its own, enable tracking (e.g. Bayley and Mogg 2023), as long as individuals are identifiable across models. Finally, if researchers wish to track animals in 2D, similar to Haalck et al. (2020) and Haalck et al. (2023), Agisoft Metashape offers the option to create geographically accurate orthomosaic projections of a 3D model (Fig.3) along an axis of choice. Metashape also enables the creation of DEMs, which could be useful for comparability to past movement or landscape ecological research.

Another limitation of our approach is that the method we have described is not well-suited for the analysis of speed or acceleration across full trajectories. This would require the plotting of points at consistent time intervals, which would result in the omission of any spatial-use details occurring between timepoints. Our approach has prioritized the estimation of 3D spatial use by moving animals, which promises to uncover characteristics of spatial use, orientation, navigation, and other as-yet unforeseen animal-habitat interactions.

Important to mention is that we did not have access to GPS devices for georeferencing at the time of our fieldwork. In the absence of such external coordinates, Agisoft Metashape produces an internal xyz-coordinate system based on the detected camera/image positions. Though we did collect depth measurements at the mantis shrimp burrows, we were not able to include these depth measurements as z-values, since Metashape automatically filled the associated x- and y-values for any given point with zeros as soon as it was assigned a z-value. If anyone’s research question should require an external coordinate system without access to GPS devices, perhaps placing perpendicular measuring tapes in the scene before collecting *Habitat Footage* would enable inclusion of xy-coordinates to complement depth/elevation measurements (z-coordinates).

The methodology we describe in this paper is enabling us to track understudied, small animals (53–153 mm long; Ahyong 2001) in the wild at an unprecedented level of detail as they traverse tens of meters of 3D reef habitat. Given the precision of photogrammetric reconstruction (e.g. Figueira et al. 2015; Guo et al. 2016), the approach we propose for plotting and measuring 3D trajectories of animals will result in impressive estimates. Further incorporating photogrammetry into studies of animal movement and sensory ecology could be immensely valuable for contextualizing animal behavior with habitat structure and externally available information.

In future, we would be excited to see the development of an automatic video-tracking software that is able to track animals through lengthy, dynamic footage and return trajectories in relative space for imposition on 3D photogrammetric models. Recently published tools that use motion to help detect and classify behaviors (e.g. Troscianko et al. 2026) hint promisingly in this direction. We furthermore believe that the fields of sensory ecology (e.g. Dangles et al. 2009; Willemart 2023) and movement ecology (e.g. Nathan et al. 2008) would benefit from a mutual exchange of tools and concepts as they transition towards three-dimensional research questions, and we encourage both fields to monitor the 3D morphometrics and visualization tools developing in landscape/seascape ecology (e.g. Vivancos, Closs, and Tentelier 2017; Kedron et al. 2019; Lepczyk et al. 2021; Yuval et al. 2023). Interdisciplinary exploration can foster greater awareness of the research questions in related fields (e.g. Demšar et al. 2015), possibly also expanding the usefulness of datasets with minimal additional effort.

## Funding

The work in this manuscript was supported by the Engineering and Physical Sciences Research Council, the International Macquarie Research Excellence Scholarship Program, the International Society for Neuroethology, The Crustacean Society, The Challenger Society for Marine Science, and the Estuarine & Coastal Sciences Association.

## Notes

### Competing Interest Statement

The authors have declared no competing interest.

